# Capturing single-cell heterogeneity via data fusion improves image-based profiling

**DOI:** 10.1101/328542

**Authors:** Mohammad H. Rohban, Shantanu Singh, Anne E. Carpenter

## Abstract

Single-cell resolution technologies warrant computational methods that capture cell heterogeneity while allowing efficient comparisons of populations. Here, we summarize cell populations by adding features’ dispersion and covariances to population averages, in the context of image-based profiling. We find that data fusion is critical for these metrics to improve results over the prior state-of-the-art, providing at least ~20% better performance in predicting a compound’s mechanism of action (MoA) and a gene’s pathway.

## Main

As a very early large-scale, high-dimensional, single-cell-resolution data type, high-throughput microscopy experiments have presented one of the first exemplars of the challenges in summarizing and comparing cell populations.

One of the key challenges is creating a *profile* of each cell population. A profile is a summary of many features of a population that enables efficient comparison with other populations while simultaneously capturing their natural variations and possible subpopulations. Recent studies have yielded many insights into cellular heterogeneity and its importance^1–4^.

Although anecdotal evidence of the value of capturing heterogeneity abounds, it has remained puzzling that so-called *average profiling*, the practice of feature-averaging all single-cell measurements together using measures-of-center (mean or median), has remained the top-ranked approach in the field of image-based (or morphological) profiling, whether those features are raw or pre-processed using unsupervised learning, or whether they derive from classical image processing or deep learning.

In image-based profiling, average profiling has been used as a straightforward way of summarizing a cell population (a sample) into a fixed length vector (a sample’s profile), with one value per feature per sample. Various metrics of similarity between profiles of two samples can then be used to infer whether they show similar phenotypic responses to their respective treatments. Average profiling typically results in a thousand-fold decrease in data size (because there are typically around a thousand cells per well in image-based profiling experiments conducted in multi-well plates), which makes downstream processing both computationally manageable and potentially statistically more robust.

However, average profiling results in loss of information about a population’s heterogeneity. The information loss can manifest in different forms. For example, multiple configurations of distinct subpopulations of cells could yield identical average profiles. Or, if two subpopulations with opposite phenotypes exist in a sample, they might cancel out yielding a profile indistinguishable from that of a sample that contains neither subpopulation. In addition to information loss, averaging can result in misleading interpretation of feature associations, e.g. Simpson’s paradox^5^. Finally, averaging makes the implicit assumption that the underlying joint features distribution is unimodal, which if violated can lead to artifacts. In this paper we investigate whether including heterogeneity measures in the profiles of cells undergoing various treatment conditions can improve upon prior methods that do not capture heterogeneity well.

Several methods have been developed in an attempt to capture cell population heterogeneity while still allowing efficient comparisons between different populations. A simple solution is to compute the cell population’s dispersion (e.g., standard deviation or median absolute deviation, MAD) for each feature and concatenate these values with the average profile. Although feature normalization brings features to comparable scales, features in average profiles generally follow a probability distribution different from that of the features in dispersion-based profiles. This discrepancy may lead to the correlation between profiles being biased toward either only features of the average or the dispersion. Concatenation can also dilute the signal-to-noise ratio (SNR) if one type of profile already has a low SNR^6^, i.e. the SNR of concatenated data would be lower than the maximum of SNRs across data types. In practice, concatenation of median and MAD profiles has been shown to provide only a minor improvement over median profiling alone^7^.

Measures of dispersion might only capture a small fraction of the heterogeneity in the data, i.e. they disregard subpopulation structures, because they involve processing each feature separately. Instead of capturing dispersion for each individual feature, one can alternatively model the heterogeneity by clustering cells using all features simultaneously. In this approach, a subset of data is used to estimate clusters of cells (representing subpopulations) and profiles are calculated as the feature averages within subpopulations^8^. Alternatively, cells can be classified into pre-determined phenotype classes using a supervised approach, and the profile is then defined as the fraction of cells in each phenotype class^9^. However, many cell phenotypes are better considered as a continuum of varying morphologies rather than discrete populations. Further, there may exist some rare subpopulations that are unique to a small portion of the data that may be overlooked in the clustering step. As a remedy, if we instead try to cluster each sample separately, the subpopulations may not be appropriately matched across samples, which makes the profiles uncomparable across the samples. Unfortunately, despite their intuitive appeal, none of these ideas have proven to significantly improve upon the baseline average profiling, at least, on the single public dataset with available ground truth, which are annotations of the mechanism of action (MOA) of a small set of compounds. In a comparison of profiling methods, average profiling (after dimensionality reduction) outperformed methods that attempted to capture heterogeneity in the data^7^. More recent work demonstrated a deep learning approach to feature extraction that yielded the highest performance yet, but nonetheless relied on average profiling^10^.

Here, we test fusing information from the dispersion profiles with the average profiles at the level of profiles’ similarity matrices. This avoids inclusion of features with inherently different probability densities in the final profile. Modeling profile similarity matrices from disparate data types using a graph has been shown to be effective in handling heterogeneous data sources such as DNA methylation, miRNA expression, and mRNA expression^6^.

We also consider alternate heterogeneity representations that do not explicitly model subpopulation information, but nevertheless capture heterogeneity. Higher order moments, which consider combinations of features, (as opposed to univariate moments of single features such as mean/median or standard deviations) are excellent candidates. As shown schematically (Figure 1), two cell populations may differ dramatically but have identical means and standard deviations. However, there is a substantial difference in the covariance (a second moment) of two features between the control (on the left) and treatment (on the right) cell populations, making this information useful to include in the populations’ profiles.

**Figure 1:**
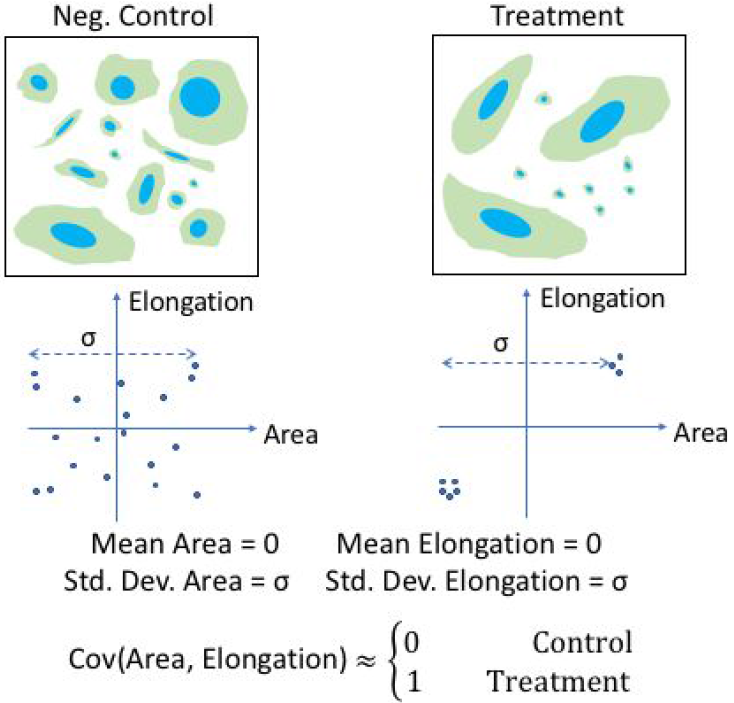
Features’ covariance captures cell phenotypes better than feature averages or dispersion, in certain situations. In this synthetic example, the negative control sample (on the left) consists of cells displaying heterogeneous morphologies. The treatment, on the other hand, shows two distinct subpopulations. In both cases, the scatter plot helps to see that the mean and standard deviation of both measured cell features (area and elongation) are equivalent in the two cases. However, the two features positively correlate in the treatment condition as opposed to the control. In such a case, the covariance can distinguish the phenotypes better than simple averages (e.g., means and medians) and measures of dispersion (e.g., standard deviations and median absolute deviations).

We also motivate the use of higher order joint statistical moments for profiling from a more theoretical standpoint. In the terminology of estimation theory, we aim to find a *sufficient statistic* for the unknown subpopulations to serve as the sample profile. A sufficient statistic is a summary of data that provides maximal information about the unknown parameter of a model that is used to explain the data. Previous work has shown that under certain assumptions, the first, second, and third order moments, collectively, are approximately a sufficient statistic for modeling subpopulations given a large enough sample size^11^ (Online Methods). Unfortunately, for typical single-cell datasets, sample sizes are too small, and computational requirements are too high, for estimating third order moments.

Here we find that even sparse random projections^12^ of covariances (second-order moments) can provide a substantial improvement in the ability to accurately compare cell populations for phenotypic similarity, when combined with median and MAD profiles via data fusion.

Testing profiling methods against each other is not a trivial exercise, given that the true similarities and differences among large sets of cell populations is rarely known. We therefore tested the approach on three different publicly available datasets where some ground truth (i.e., expected results), albeit imperfect, is known. Cell measurements in the datasets are based on Cell Painting, which is an image-based assay designed to capture cell morphology^13^. For these benchmark datasets, our laboratory had released the image data^14,15^ but for this study we collected ground truth to create a proper testing scenario. We used datasets that had a sufficient number of perturbations for the data fusion technique to work (Online Methods), and therefore did not include the dataset reported in a previously published study^7^.

To evaluate each profiling method, we tested whether pairs of cell populations that look most alike, according to the computed image-based profiles, have been treated with perturbations that are annotated as having the same mechanism of action (for compounds) or the same pathway (for gene overexpressions). Similarity between pairs of image-based profiles are established based on the profiles’ correlation (Online Methods).

We find that the enrichment of top-correlated perturbation pairs, whether genetic or chemical, in having same mechanism of action or same pathway (Online Methods), is improved when median absolute deviation and/or covariances (summarized by sparse random projections to avoid the curse of dimensionality) are combined with the median profiles through Similarity Network Fusion (SNF) (Online Methods) (Figure 2A and Supp. Tables 1 and 2). The improved intra-MOA similarities, especially in certain MOAs (Figure 2B, Supp. Tables 3–5), indicates that median, MAD, and sparse projections of covariances are complementary sources of information.

The improvement in enrichment of top-correlated perturbation pairs is observed across the three datasets, which involve different experimental conditions and perturbation/annotation types. Although trivial concatenation of dispersion measures with the median profile has shown marginal improvement in the past^7^, combining MAD and covariance with medians via data fusion provides consistent and substantial improvement (typically around 20%) in the mentioned enrichment score. We conclude that capturing cell-to-cell heterogeneity is of value in image-based profiling of cell populations. Source code, image processing pipelines, and gene/compound annotation data to reproduce and build on these results are available (https://github.com/carpenterlab/2018rohbansubmitted).

**Figure 2:**
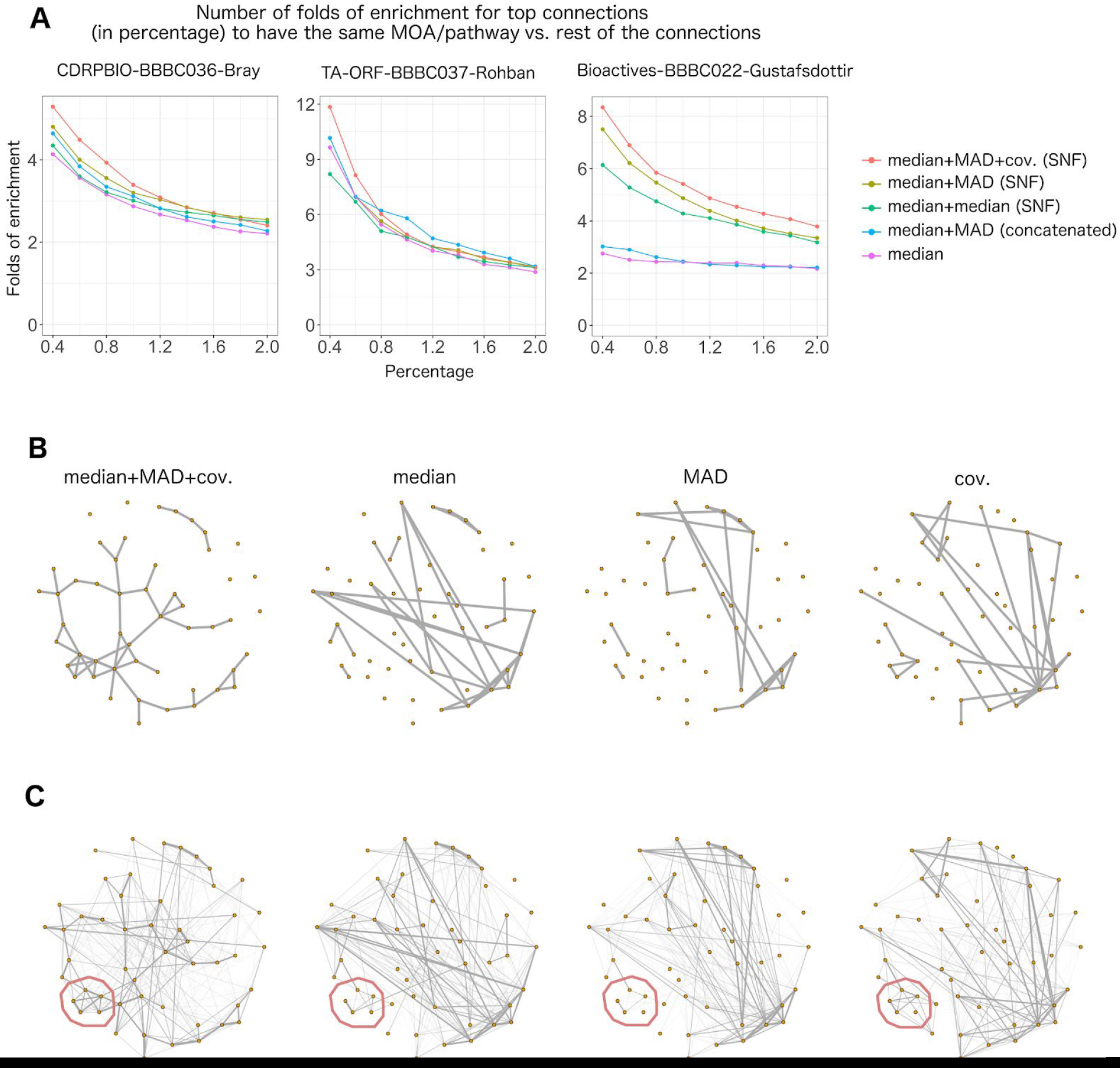
Incorporating metrics of cell heterogeneity via data fusion increases the percentage of connections between image-based profiles that are validated. **A:** When median, MAD and random projections of covariance profiles are combined through SNF (red line), the enrichment in having same MOA/pathway annotations is improved, especially for the strongest, most relevant connections above 0.5%. This is shown in three separate experiments involving small molecules (left, right) and gene overexpression (middle). Enrichment is versus a null distribution, which is based on the remainder of the connections. **B:** Similarity graphs for the mechanism-of-action (MOA) class “Adrenergic receptor antagonists”, using different types of profiles in CDRPBIO-BBBC036-Bray. This MOA was chosen because it showed the highest improvement upon combining different profiles. Each node represents a compound, and two nodes are connected if the similarity of their corresponding profiles is ranked among the top 5% most-similar pairs. Median, MAD, and random projections of covariance profiles seem to be complementary for this MOA, as they cover mostly non-overlapping compound connections. The overall connectivity of compounds in this MOA is improved once these profiles are combined through SNF. Graph layouts are the same across data types and are based on the similarities in median+MAD+cov. (SNF). **C:** Weighted similarity graph as in the previous plot except that edge thicknesses are based on an exponential weighting of the ranked similarity values. Sub-clusters that are moderately present in two or three profile types (such as the one marked in red on bottom left) became stronger after applying data fusion using SNF.

## Online methods

Source code, image processing pipelines, and gene/compound annotation data to reproduce and build on these results are available (https://github.com/carpenterlab/2018_rohban_submitted).

### Datasets

We used three datasets to evaluate the profiling methods:

- *CDRPBIO-BBBC036-Bray*: 2200 known bioactive compounds in U2OS cells. This dataset is the bioactive subset of a publicly available dataset^15^. Raw images are available at https://idr.openmicroscopy.org/webclient/?show=screen-1251.
- *Bioactives-BBBC022-Gustafsdottir*: 1600 known bioactive compounds in U2OS cells. This is the image set BBBC022v1^14^, available from the Broad Bioimage Benchmark Collection^7^. Raw images are available at https://data.broadinstitute.org/bbbc/BBBC022 and https://idr.openmicroscopy.org/webclient/?show=screen-1952. The compounds in this dataset have some overlap with CDRPBIO.
- *TA-ORF-BBBC037-Rohban*: ~200 genes in various signaling pathways are over-expressed in U2OS cells^16^. Raw images are publicly available at https://idr.openmicroscopy.org/webclient/?show=screen-1751.

In all three datasets, around 1700 single cell image-based readouts were obtained by running the Cell Painting assay^13^ and an image processing pipeline in CellProfiler^17^ software. The features are z-scored platewise in all datasets based on the negative controls.

Extracted image-based features are publicly available in the following s3 bucket s3://cellpainting-datasets under folders corresponding to the respective names of the datasets.

### Annotations

We used the Repurposing hub^18^, https://clue.io/repurposing-app, to annotate compounds with their mechanism-of-action (accessed on Feb. 28th 2018). For missing annotations, we used other resources such as https://www.drugbank.ca. The gene overexpression dataset contains biological pathway annotations, generated by domain experts at our institution. For pathways marked as having “canonical” and “non-canonical” members, we merged all members.

### Theoretical justification for using moments to describe cell populations

The theoretical basis we present assumes that the cellular phenotypes can be modelled as a mixture of gaussians. This model has been shown to be effective in capturing subpopulations in imaging data^8,19^. In this model, the subpopulations and their proportions correspond to mixture centers and mixture prior probabilities, respectively. Both of these quantities are considered as unknown parameters.

It has been shown that, under mild assumptions, these unknown parameters can be estimated using the first, second, and third moments of data^11^. More specifically, if the mixtures are spherical Gaussians, and their centers are linearly independent, all the unknown parameters can be estimated with high precision given a sufficiently large number of data points (see Theorems 2 and 3^11^). In other words, the first, second, and third moments of the data constitute an approximate sufficient statistic for the unknown parameters in GMM when the sample size is sufficiently large. Average profiling uses only a small portion of this sufficient statistic-the first moment-to represent the sample. We can make this representation richer by also including the second and third moment profiles. Going beyond the third moment does not add any additional information with regard to the GMM (Theorems 2 and 3^11^).

We did not test the third moment because our datasets contain in the order of a few thousand cells per sample, whereas millions of cells would be needed to robustly compute third moments (O(d^3^), where d is dimensionality of the feature space; on the order of 100 in our case). As well, the dimensionality of the final profiles rapidly grows as d increases. Although computing second-order moments is more feasible, it nonetheless requires dimensionality reduction to be of practical use: for 500 features, the second moment is nearly 125,000-dimensional, which is both computationally and statistically difficult to work with in forming the treatments similarity matrix. We use sparse random projections^12^ of the vectorized covariance matrices to reduce the dimensionality to 3000 covariances while approximately preserving pairwise profile distances.

### Combining first and second order moments

Because the statistical distributions of mean, MAD, and covariance profiles can be different in general (Supp. Figure 1), we combined them at the sample similarities level, rather than simply concatenating the profiles. We use Similarity Network Fusion (SNF)^6^, which operates on a graph representation of the dataset in each data type (in our case, three: medians, MADs, and covariances). A graph diffusion process then is used to combine the graph for each data type into the final network, which encodes the pairwise similarity values. SNF has shown great promise in fusing biological readouts when the number of samples is on the order of few hundreds^6^. The method is expected to be less effective when sample size falls below a threshold as local neighbors start to become less similar, violating the assumptions made in the method^20^. For this reason, we did not use the prior benchmark dataset in^7^.

### Parameter settings

We used the *SNFtool* R package (Ver. 2.2.1) for data fusion to combine data types (median, MAD, and covariance profiles), and set the neighborhood size k = 7 in the similarity graph, gaussian weight function bandwidth μ = 0.5, and number of iterations T = 10 (for two data types) and T = 15 (for three data types) in SNF, which are typical choices for the algorithm. To avoid overfitting, we did not test alternative values of these parameters. Prior to applying SNF, similarity matrices are z-scored based on median and MAD and then linearly scaled to map 99.9th percentile to 0.999. This helps to make sure that the similarity values are on the same scale across data types.

We used 3000 sparse random projections with the density of p = 0.1 (the probability of an entry in the random projection matrix of being non-zero) to reduce the dimensionality of the covariance profiles in all the datasets. We observed reasonable consistency against randomness in the treatment correlation matrices when using around 3000 random projections (Supp. Figure 2). Pearson correlation of profiles is used to form similarity matrices, which are used as the inputs to SNF.

### Evaluation tasks

We evaluated different profiling strategies in this paper (Figure 2) based on whether the most-similar treatment pairs (above a given cutoff) are enriched for having the same MOA/pathway annotation, after removing un-annotated compounds. To ensure that strong profile similarities are not driven by systematic effects that might make samples on the same plate look more similar to each other than to those in other plates, all same-plate pairs were excluded in this analysis.

We rejected an alternative evaluation approach, accuracy in MOA/pathway classification^7^, which only works well if MOAs are all well/equally represented in the dataset. The approach we took is better suited for the MOA class imbalance situation (as is the case for the datasets analyzed in this paper), as the enrichment is calculated based on a null distribution that tends to normalize MOA class sizes implicitly. Otherwise, treatments belonging to larger MOA classes tend to dominate the classification accuracy.

### Enrichment score

We define enrichment score as the odds ratio in a one-sided Fisher’s exact test, which tests whether having high profile similarity for a treatment pair is independent of the treatments sharing an MOA/pathway. To perform the test, we form the 2×2 contingency table by dividing treatment pairs into four categories, based on whether they have high profile correlations, determined by a specified threshold (in rows) and whether they share an MOA/pathway (in columns). The odds ratio is then defined as the ratio of elements in the first row divided by that of second row in the contingency table. This roughly measures how likely it is to observe same MOA/pathway treatment pairs in highly correlated vs. non-highly correlated treatment pairs.

## Supplementary Figures and Tables

**Supplementary Table 1:** The proposed median+MAD+covariance (data-fused) profiles significantly improve *recall* vs. state-of-the-art median+MAD (concatenated). More specifically, the percentage of same MOA/pathway connections that are captured in the top 0.5% most-similar treatment pairs is significantly higher in median+MAD+covariance (data-fused) profiles compared to median+MAD (concatenated) profiles in CDRPBIO-BBBC036-Bray and Bioactives-BBBC022-Gustafsdottir (Fisher’s test). The number of genes or compounds in the experiment is indicated by “n” in the table.

| Dataset | Relative increase in validated hits | P-value (Fisher's test) |
| --- | --- | --- |
| CDRPBIO-BBBC036-Bray<br>(n compounds = 1552) | 19% | 0.038 |
| Bioactives-BBBC022-Gustafsdottir<br>(n compounds = 1048) | 132% | 1.227e-11 |
| TA-ORF-BBBC037-Rohban<br>(n genes = 205) | 12% | 0.36 |

**Supplementary Table 2:** The proposed median+MAD+covariance (data-fused) profiles significantly improve *precision* vs. state-of-the-art median+MAD (concatenated). More specifically, the top 0.5% most-similar treatment pairs contain more same MOA/pathway pairs in median+MAD+covariance (data-fused) profiles compared to median+MAD (concatenated) profiles in CDRPBIO-BBBC036-Bray and Bioactives-BBBC022-Gustafsdottir (Fisher’s test). The number of genes or compounds in the experiment is indicated by “n” in the table.

| Dataset | Relative increase in validated hits | P-value (Fisher's test) |
| --- | --- | --- |
| CDRPBIO-BBBC036-Bray<br>(n compounds = 1552) | 20% | 0.036 |
| Bioactives-BBBC022-Gustafsdottir<br>(n compounds = 1048) | 140% | 4.458e-12 |
| TA-ORF-BBBC037-Rohban<br>(n genes = 205) | 18% | 0.33 |

**Supplementary Table 3:**
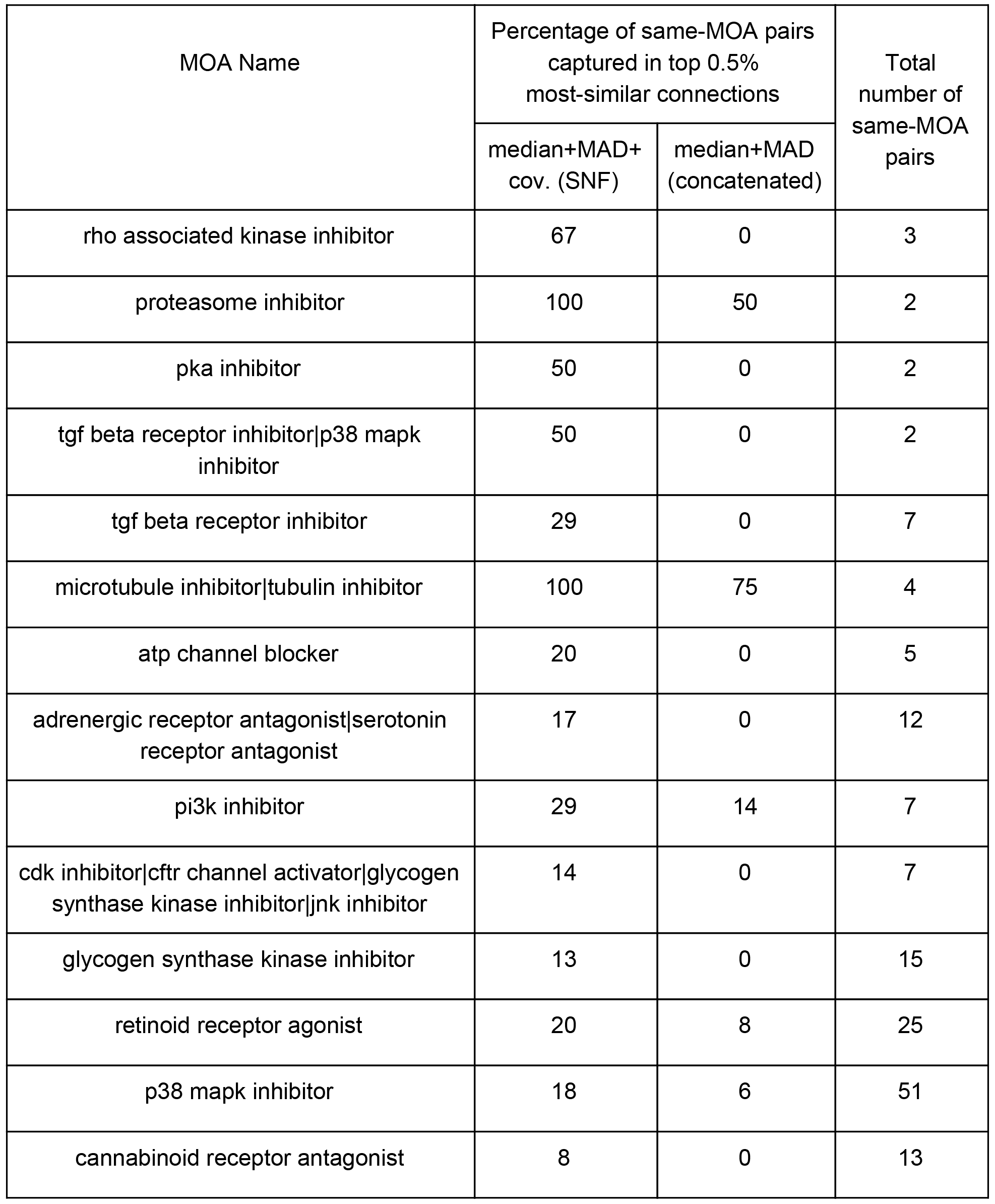
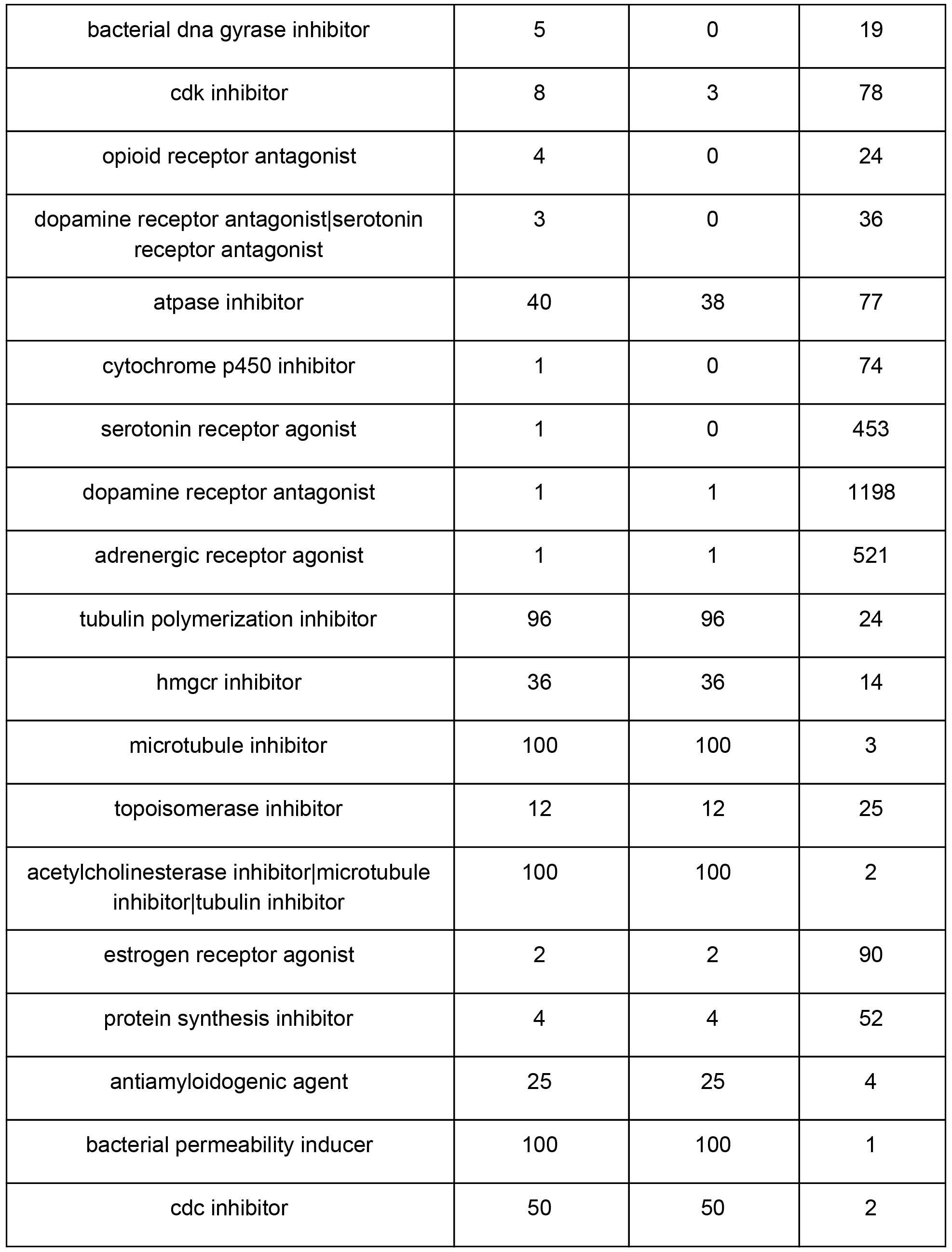
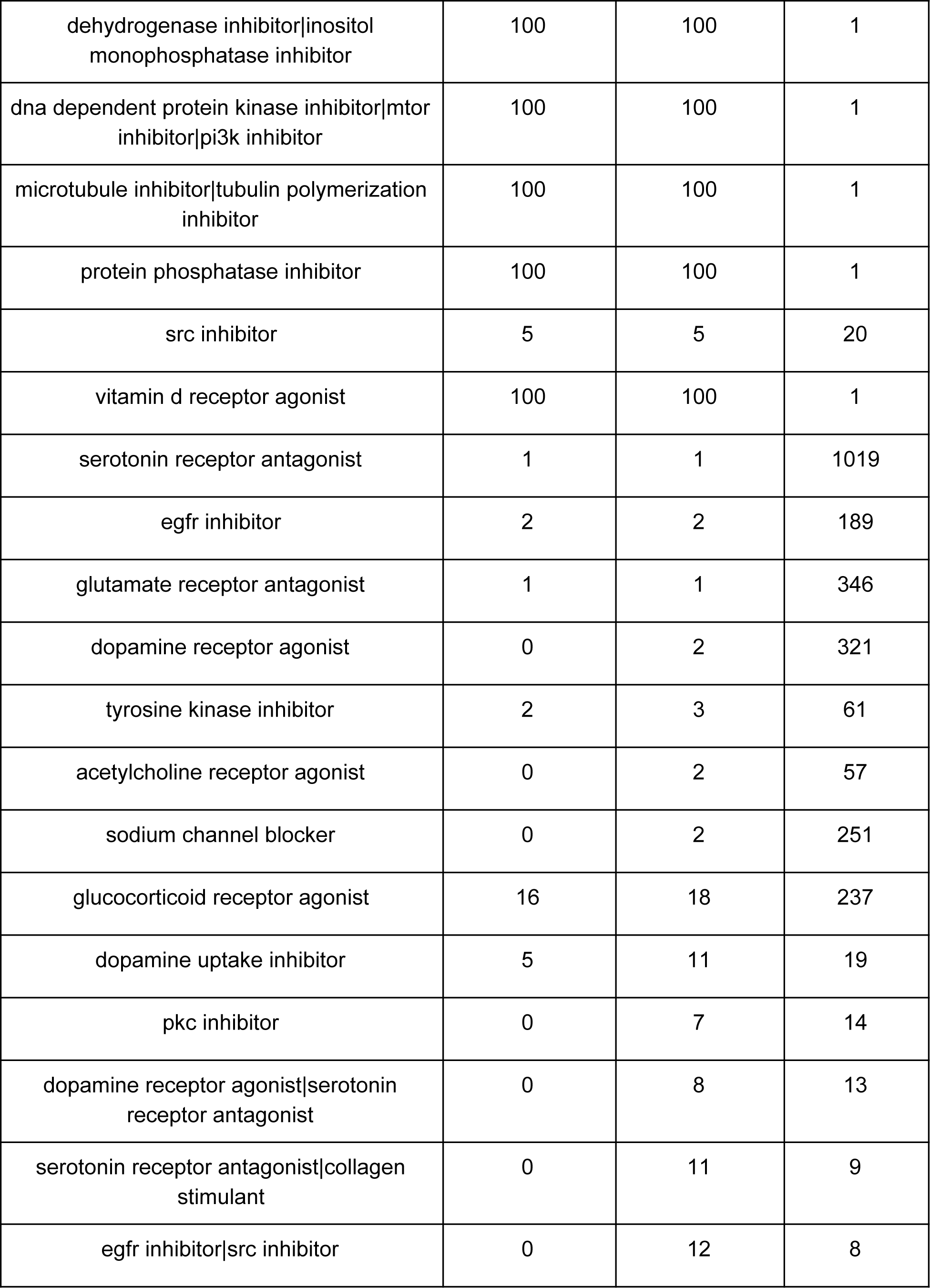
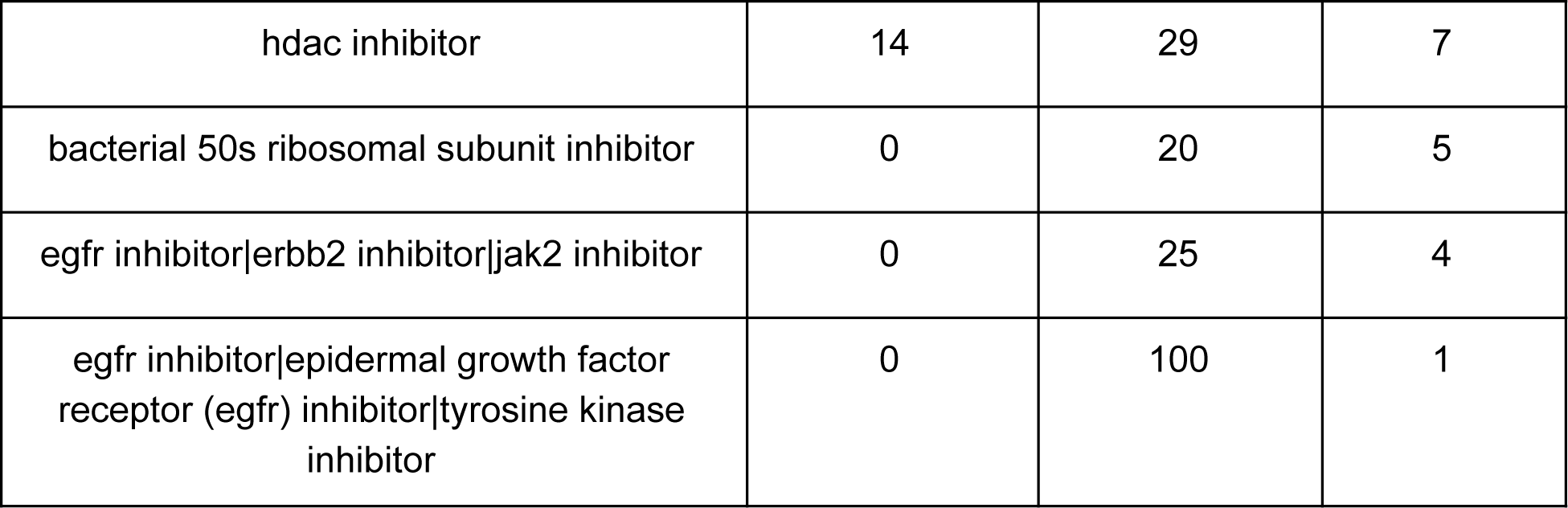
Sorted list of MOAs based on improvement of median+MAD+cov. (SNF) compared to state-of-the-art median+MAD (concatenated) in CDRPBIO-BBBC036-Bray. TGF-beta receptor inhibitors, ATP channel blockers, Tubulin inhibitors, and Glycogen synthase kinase inhibitors are among the MOAs showing improvements.

**Supplementary Table 4:**
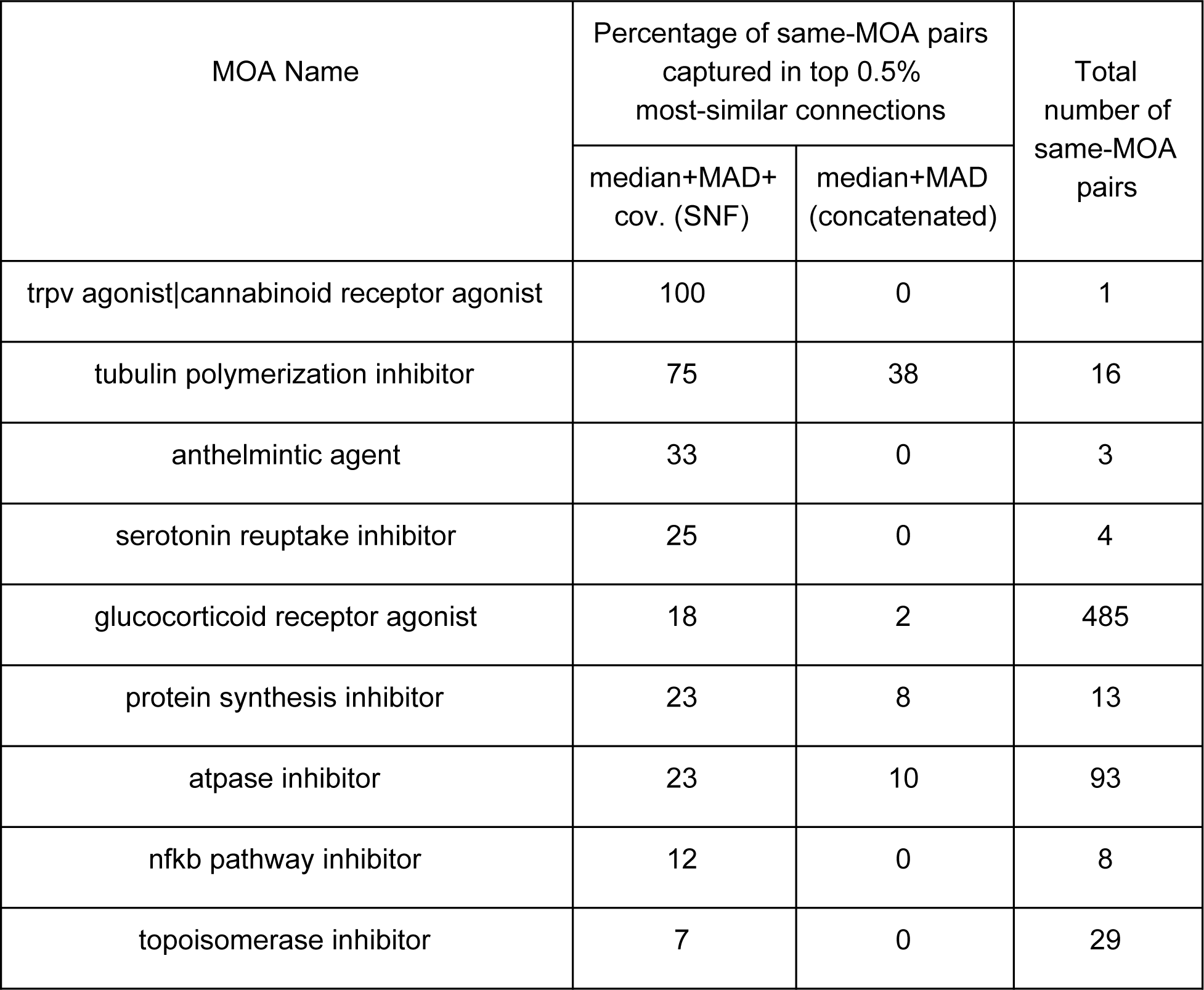
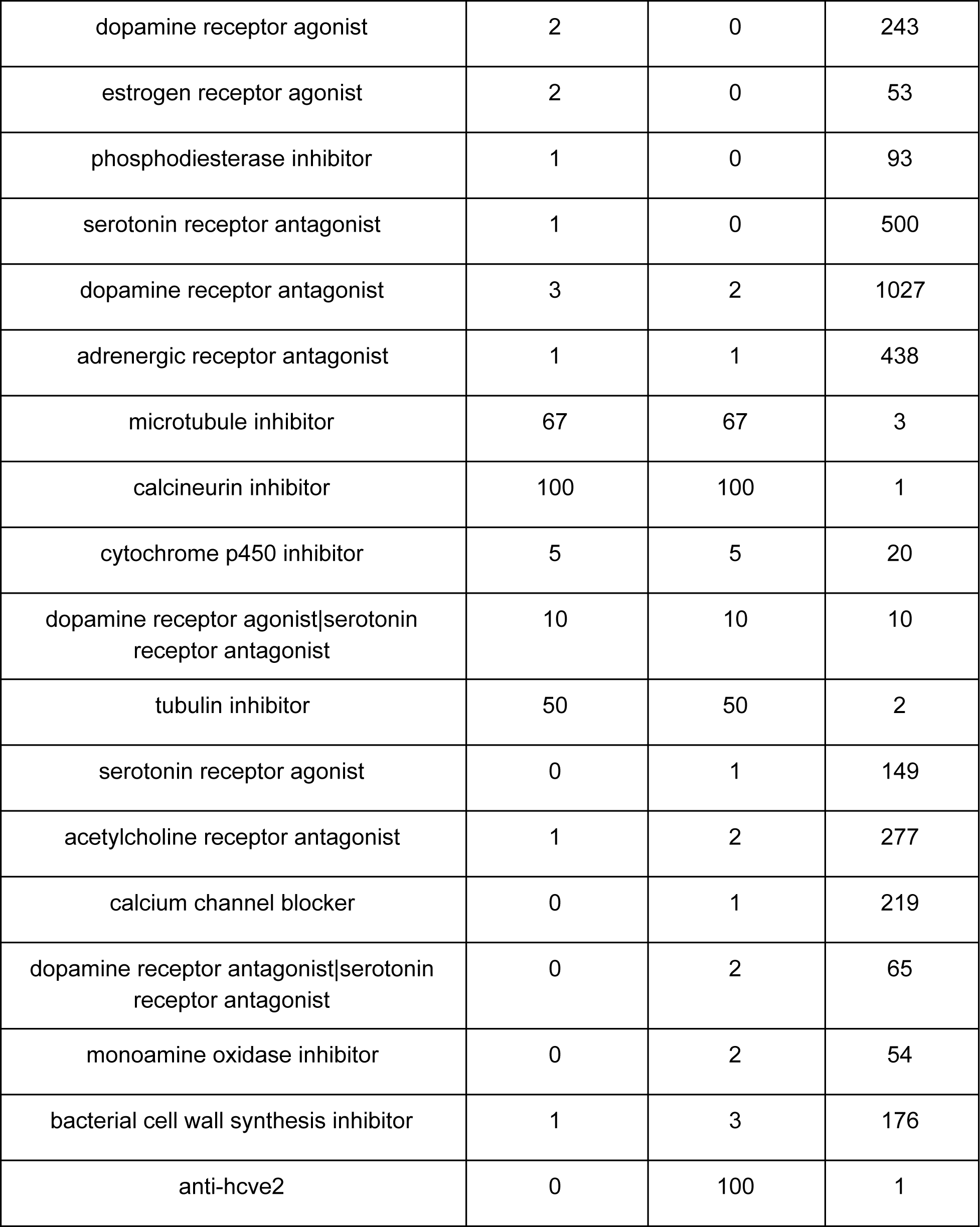
Sorted list of MOAs based on improvement of median+MAD+cov. (SNF) compared to state-of-the-art median+MAD (concatenated) in Bioactives-BBBC022-Gustafsdottir. Tubulin inhibitors and Glucocorticoid receptor agonists are among the MOAs showing improvements.

**Supplementary Table 5:**
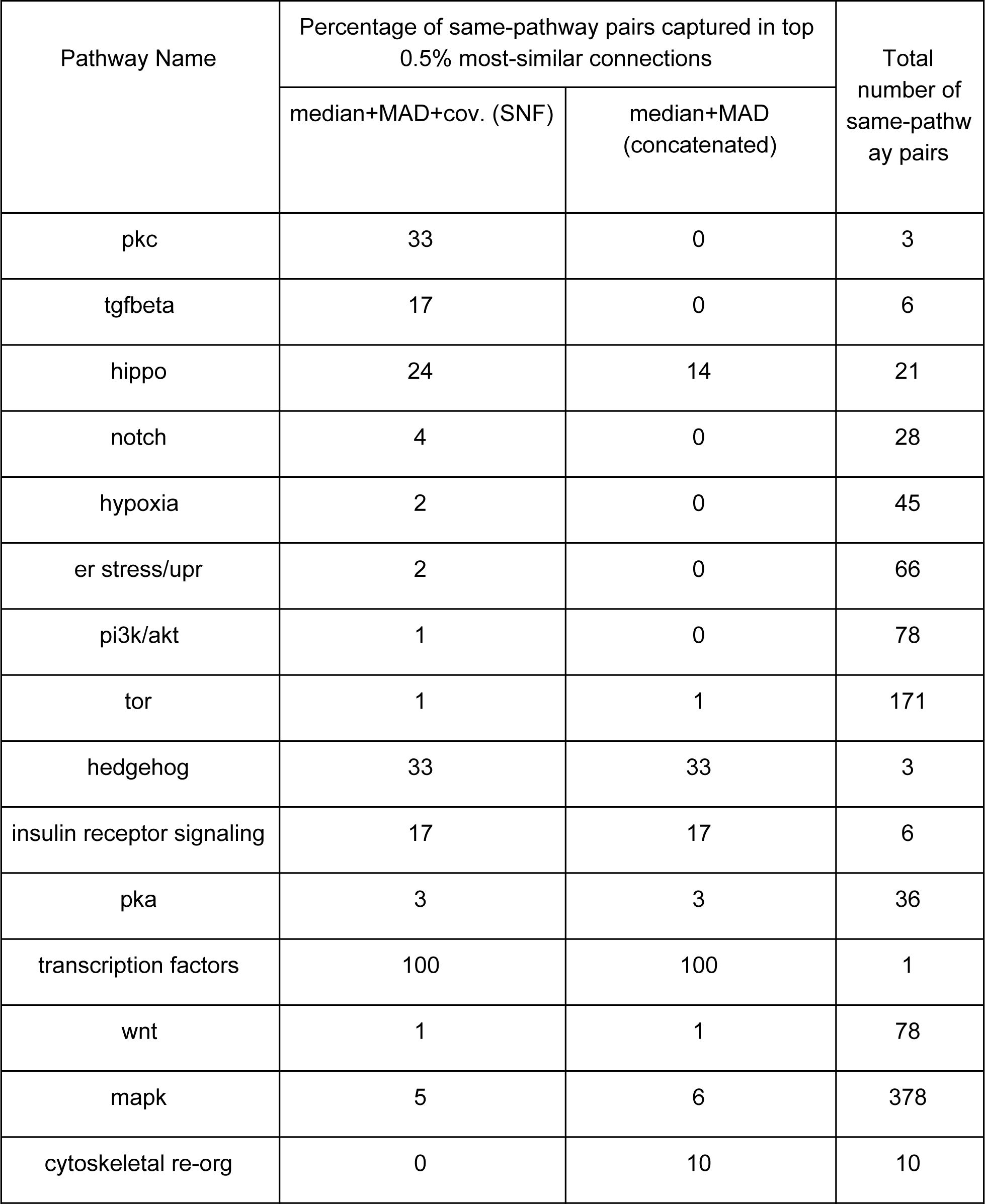
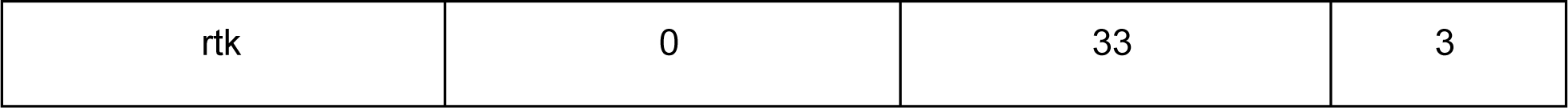
Sorted list of pathways based on improvement of median+MAD+cov. (SNF) compared to state-of-the-art median+MAD (concatenated) in TA-ORF-BBBC037-Rohban. PKC, TGF-beta, Hippo, and NOTCH are among the pathways showing improvements.

**Supplementary Figure 1:**
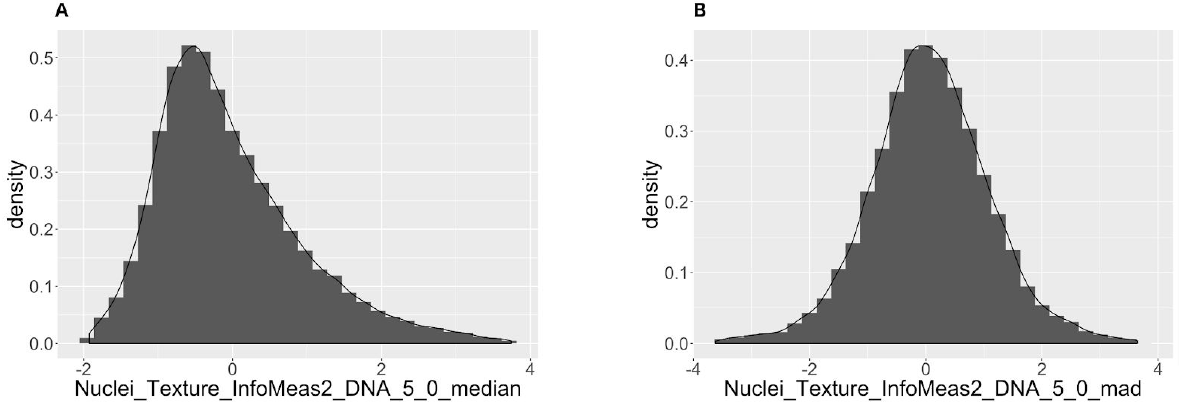
Features may show different distributions in median and MAD profiles. (A) shows that a texture feature in the DNA channel in median profiles has a skewed distribution in CDRP-BBBC036-Bray. On the other hand, MAD profiles gives a nearly symmetric distribution for the same feature (B). Values lower than first and higher than 99th percentiles are removed before plotting the distributions.

**Supplementary Figure 2:**
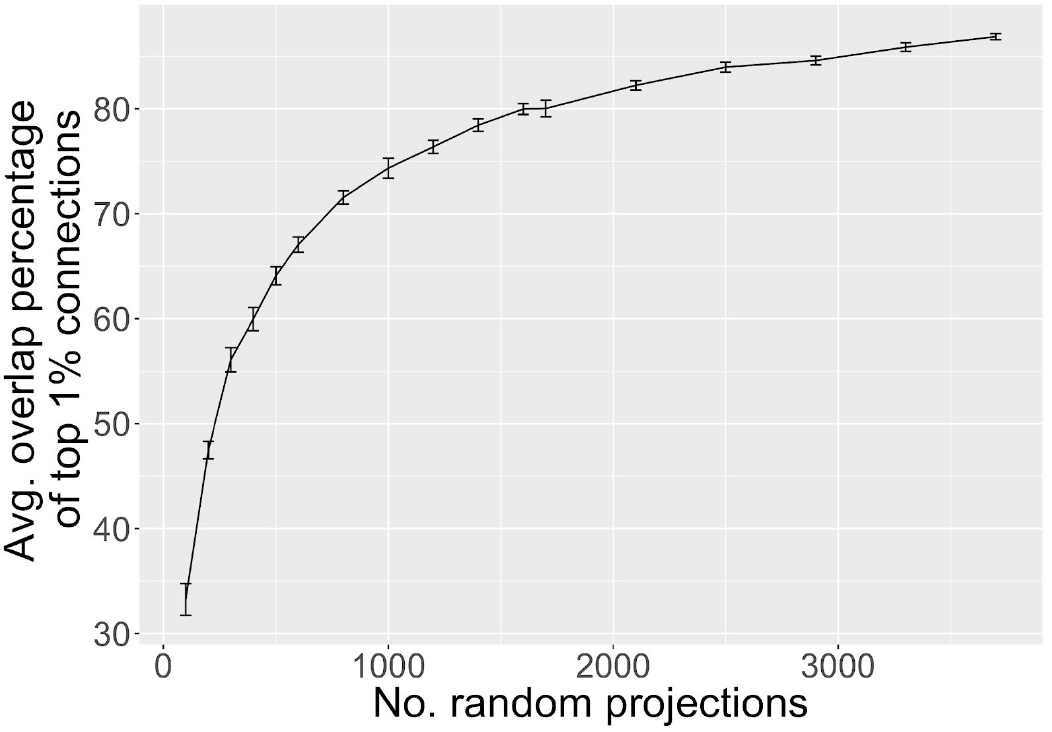
Top-correlated treatment pairs become increasingly consistent as number of random projections in increased. Average percentage of overlap size between top 1% correlated treatment pairs between two random projections increases sharply and saturates around 3000 random projections. The data is taken from a single plate in the CDRP-BBBC036-Bray dataset and each point in the plot is the average of 20 independent random simulations.

